# A Mammalian Surface Display Platform to Optimize the Antigenicity of Viral Proteins for Vaccine Design

**DOI:** 10.64898/2026.01.09.698728

**Authors:** Jayani Christopher, Caitlin Harris, Daniel J. Marston, Brianna Rhodes, McKenzie Frazier, Mihai L. Azoitei

## Abstract

Vaccine development often involves modifying native viral proteins to enhance their stability and antigenicity, as seen in approved Covid-19 vaccines and multiple HIV vaccine candidates currently under investigation. High throughput screening on the surface of mammalian cells enables the rapid evaluation of oligomeric, glycosylated viral proteins and the identification of mutations that improve their properties for vaccine design. Here, we developed an experimental platform that uses the *PiggyBac* transposon system to display libraries of viral protein variants on the surface of mammalian cells for efficient screening. This approach addresses common challenges in existing mammalian display systems, including low transfection efficiency and the development of stable cell lines for iterative selection rounds. The new platform was validated by expressing and characterizing antigenically diverse viral proteins from influenza, SARS-CoV-2, and HIV. In a further application, library screening of influenza haemagglutinin libraries identified mutations that increased binding of broadly cross-reactive antibodies to a conserved, but partially occluded, epitope of interest for the development of a universal influenza vaccine. These results demonstrate the potential of this mammalian display platform to rapidly engineer immunogens with desired antigenic properties for vaccine design.

## Introduction

Rational vaccine development strategies typically employ structure-guided design and high-throughput library screening to optimize viral antigens for enhanced immunogenicity and stability. Structural analysis can guide the rational modification of natural viral proteins to stabilize preferred conformations and to expose epitopes for optimal immune recognition.^1-6^ For example, most Covid-19 vaccines employ spike proteins stabilized in the prefusion conformation through two proline substitutions (Spike-2P),^7^ a modification that greatly enhances immunogenicity. Immunogens derived from the Envelope (Env) trimer of HIV-1 are typically stabilized by adding disulfide bonds linking its subunits and by incorporating targeted point mutations to increase the solubility and stability of the native viral protein.^1, 2, 8-11^ High throughput library screening approaches can facilitate rational immunogen design by rapidly evaluating thousands to millions of candidates to identify those with desired antigenic properties.^12-21^

Multiple high-throughput experimental platforms have been developed and applied to optimize the expression, stability, or antigenicity of viral proteins. Yeast surface display has been effective for screening and engineering monomeric proteins, like the RBD domain of the SARS-CoV-2 spike or the gp120 of the HIV Env protein. However, yeast display systems are limited by their inability to display multimeric proteins and by the post-translational glycosylation modifications they generate, which are significantly different from those in mammalian cells. ^22-24^ This limits the development of virus-derived antigens that are typically highly glycosylated oligomeric proteins, as observed for coronaviruses, HIV, influenza, and others. To address this, mammalian surface display platforms enable the expression and evaluation of viral glycoproteins in their close to native, post-translationally modified forms. ^12, 18, 19, 25-27^ Based on their transfection method, current mammalian display methods can be divided into two classes, depending on whether they rely on transient transfection or lentiviral transduction, each with its advantages and weaknesses. Transient transfection is technically straightforward and enables rapid expression of variant libraries, but it does not generate stable cell line libraries that can be interrogated over multiple selection rounds. ^19, 25, 26^ Lentiviral approaches, in contrast, support stable genomic integration and allow for iterative sorting cycles, but the workflow is technically demanding, time consuming, and carries potential biosafety concerns associated with handling and producing viral vectors.^18, 21, 28^ Vector construction, viral packaging, titer optimization, and the establishment of stable clones often require several weeks to months, and these steps may still yield viral titers insufficient for robust expression.^29^ Other non-viral strategies for achieving stable expression have been described, but these methods are laborious and involve numerous sequential experimental steps.^12, 30^ Consequently, there is a need for a system that enables rapid transfection while maintaining stable and long-lasting gene expression for iterative library screening.

To address this, here we developed a mammalian surface display platform that employs the *PiggyBac* transposon machinery for stable integration and controlled expression of protein libraries. *PiggyBac* serves as a non-viral transposon system capable of efficiently integrating exogenous DNA into the host genome.^31, 32^ Unlike lentiviral workflows, *PiggyBac* does not require viral packaging or specialized biosafety procedures, and its setup is rapid and straightforward, resembling the simplicity of transient transfections. As a result, *PiggyBac* effectively combines the advantages of both transient and lentiviral methods by offering a short, less labor-intensive library transfection process while still enabling stable genomic integration, which is essential for selection across multiple sequential rounds. Previous studies have demonstrated the use of *PiggyBac* transposition to express full length IgG^33^ and single-chain Fab^34^ in mammalian surface display platforms, highlighting its utility for antibody engineering. However, these earlier systems relied primarily on constitutive promoters and lacked inducible control over protein expression, which can limit their adaptability for applications requiring regulated surface display. The mammalian surface display platform described here was validated using multiple viral membrane proteins, including influenza Hemagglutinin (HA), HIV Env, and SARS-CoV-2 spike, confirming surface expression and antigenic integrity. We further applied this system to identify mutations in the influenza HA head domain that expose the 220-loop to increase its recognition by target antibodies. This conserved, immunologically subdominant site is partially occluded at the interface of adjacent HA head domains in the native trimer, which likely limits its accessibility during natural infection or vaccination and contributes to the low frequency of antibody responses such as FluA-20^35^ and S5V2-29.^36^ These antibodies which cross react with almost all influenza A viruses target the 220-loop and are therefore of significant interest for universal influenza vaccine design. We hypothesized that engineering HA variants with increased presentation of this partially hidden epitope would enhance antibody recognition and could promote the elicitation of similar broadly reactive responses upon vaccination. This mammalian display platform can therefore facilitate the rapid engineering of viral proteins with desired antigenic profiles that are of interest for vaccine development.

## Results

### Plasmid design

To develop the mammalian display platform, we used the trimeric HA protein that decorates influenza virions as a test case to determine the optimal experimental conditions. The ectodomain residues 17–528 of the A/H3N2/Hong Kong/1968 strain were linked to a CD5 leader peptide at the 5′ end, which directs the protein to the secretory pathway, and to a c-Myc tag followed by the PGDFR transmembrane domain at the 3′ end.^19, 26^ In this configuration, the expressed HA trimer is secreted for cell surface display and anchored on the cell membrane **(Figure 1**). This DNA construct was ligated at the NotI and NheI restriction sites into a commercially available *PiggyBac* transposon vector (System Biosciences), where it was flanked on both sides by inverted terminal repeats (ITRs) to allow transposition (**Figure 1**). This vector contains a puromycin resistance gene to efficiently select the transfected cells and a cumate inducible switch regulated by the EF1-CymR repressor to control the initiation of HA expression.^37^ HEK293T cells were co-transfected with the transposon plasmid containing HA ectodomain and a *PiggyBac* transposase expression vector (System Biosciences). Once displayed on the cell surface, HA trimer expression was detected with an anti-c-Myc antibody labeled with fluorescein isothiocyanate (FITC). To ensure proper folding, known antibodies, FluA-20,^35^ S5V2-29,^36^ 429B01,^38^ and F045-092^39^ were bound to HA and labeled with an anti-human IgG monoclonal antibody conjugated with phycoerythrin (PE) fluorophore. Subsequent analysis of fluorescently labeled cells by Fluorescence Activated Cell Sorting (FACS) provided a two-dimensional analysis of individual cells for surface expression and target antibody binding (**Figure 1**).

**Figure 1.**
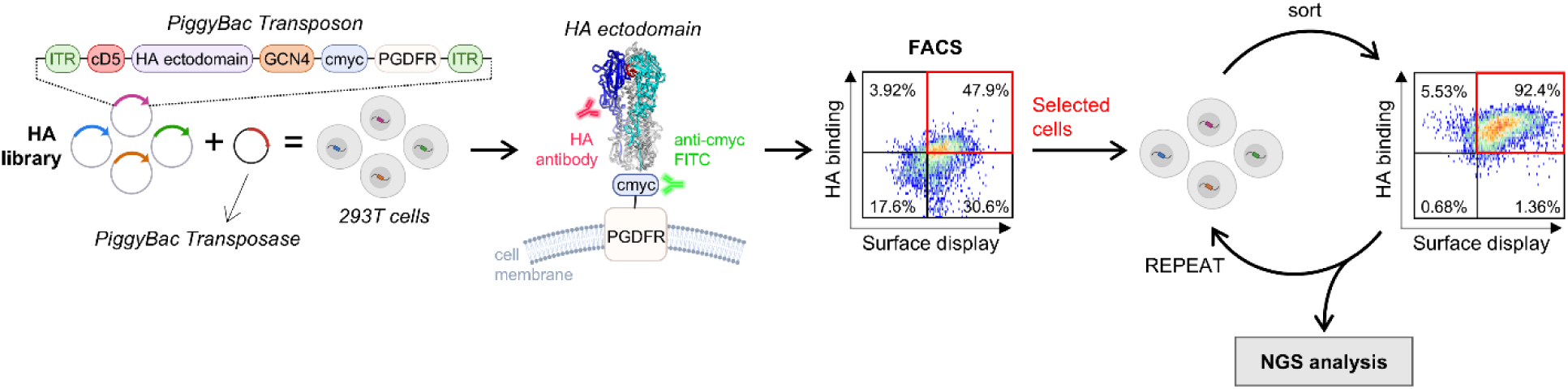
Overview of the PiggyBac-based mammalian surface display platform. The viral protein of interest is fused to a c-Myc–tagged platelet-derived growth factor receptor (PDGFR) transmembrane domain to enable anchoring on the cell surface, and the expression cassette is flanked by inverted terminal repeats (ITR). Upon induction, the protein is displayed on the cell surface and labeled with a target antibody and an anti–c-Myc FITC-conjugated antibody. Iterative rounds of FACS are performed to enrich binding populations, followed by NGS to analyze selected variants

To improve the stability and surface expression of HA trimer in mammalian cells we fused a GCN4 trimerization motif at the N-terminus of the HA ectodomain sequence. The GCN4 motif is a well characterized leucine zipper derived coiled-coil domain that promotes stable trimer formation through parallel α-helical interactions and has been widely used to stabilize oligomeric viral glycoproteins.^40^ This motif was intended to promote trimer formation to capture the native oligomeric structure of HA. Flow cytometry analysis showed a significant improvement in HA surface expression in cells transfected with the construct fused to GCN4 compared to those lacking the oligomerization motif. A higher percentage of transfected cells exhibited target antibody binding when the GCN4 trimerization domain was present. The trimerization motif therefore stabilizes HA trimers and increases their expression on the mammalian cell surface (Figure 2c). The transposon vector also contained an internal ribosome entry site (IRES) followed by a Green Fluorescence Protein (GFP) gene between the ITRs, which was present in the commercial transposon backbone used in this study. The IRES sequence allows co-expression of HA and GFP from a single transcript with GFP acting as a cytoplasmic reporter. Inclusion of GFP allowed us to easily monitor the stable integration and expression levels of the transfected construct by microscopy. Interestingly, the co-expression of GFP also increased HA surface expression as shown by FACS analysis (Figure 2d).

**Figure 2.**
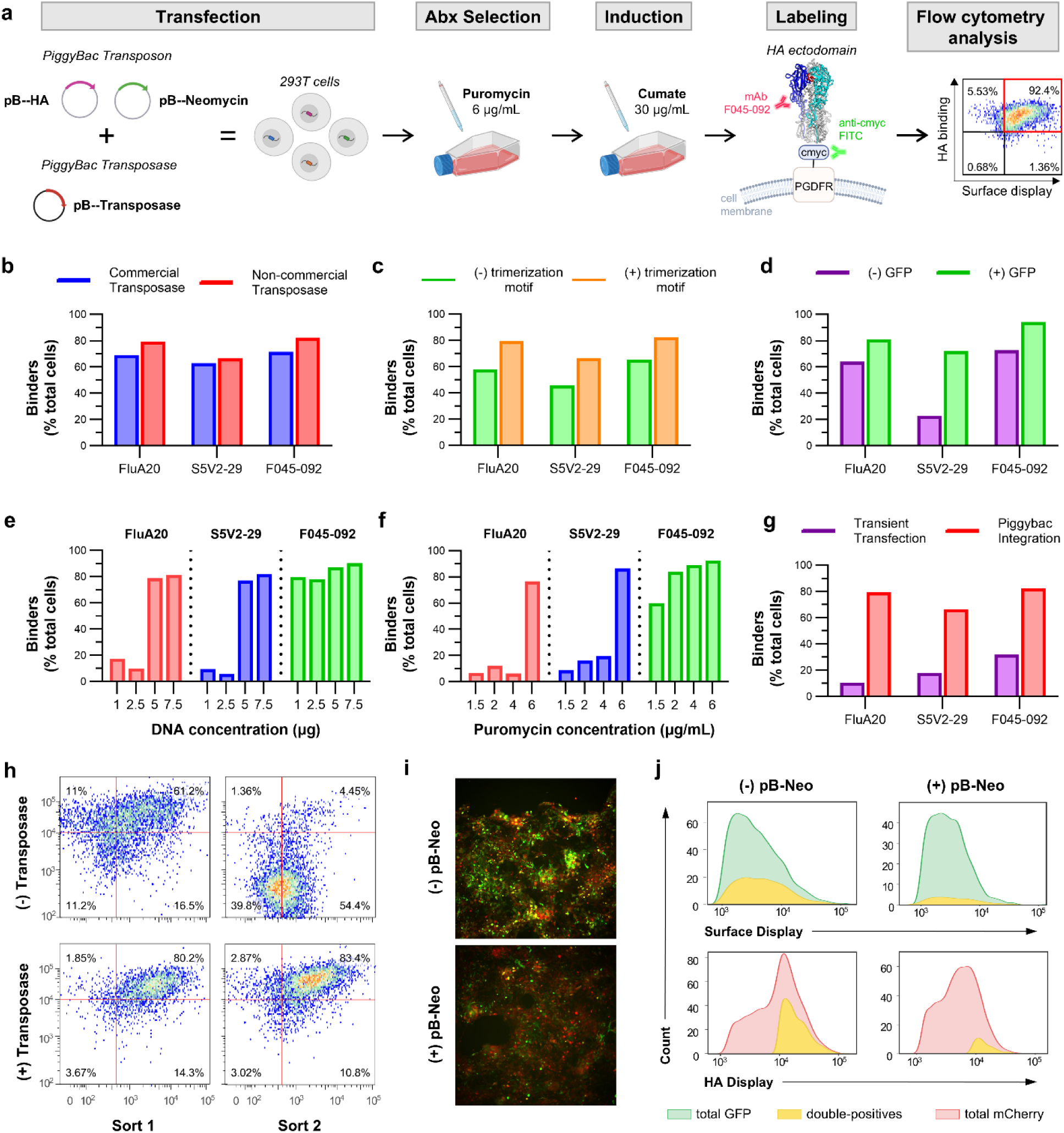
Optimization and validation of the mammalian surface display platform. (a) Overview of the optimized transfection workflow and downstream analysis, with key experimental parameters indicated. (b) Comparison of HA surface expression upon transfection and transposition using a commercial Super *PiggyBac* transposase or a non-commercial hyperactive *PiggyBac* transposase. Binders are defined as cells displaying surface-expressed antigen and simultaneously binding target antibodies, identified by FACS as double-positive populations labeled with anti–c-Myc FITC (to detect surface expression) and anti-human IgG PE (to detect bound antibody). (c) Effect of incorporating a GCN4 trimerization motif on HA surface expression. (d) Influence of GFP co-expression on HA surface expression (e) Optimization of total DNA input and (f) puromycin selection concentration for maximal HA surface expression. (g) Comparison of HA surface expression following *PiggyBac* mediated genomic integration versus transient transfection using a non-*PiggyBac* vector (h) Analysis of HA expression stability following cell sorting in the presence or absence of transposase. (i) Optimization of single-copy integration by diluting target vectors with a carrier *PiggyBac* transposon. Transfected cells are visualized by fluorescence microscopy. (j) Flow cytometry analysis of GFP and mCherry tagged HA to identify double-positive cell populations following dilution with the carrier transposon.

For stable integration, we initially co-transfected the HA containing plasmid with a commercially available vector that encodes the Super *PiggyBac* transposase (System Biosciences). This PiggyBac transposase is optimized for high expression, stability, and integration activity in mammalian cells.^37^ However, the commercially available plasmid cannot be propagated in *E. coli*, increasing experimental costs. To identify a more cost-effective alternative, we tested an engineered hyperactive PiggyBac transposase^41^, which binds to the same ITR regions as the Super PiggyBac transposase. We co-transfected the engineered hyperactive PiggyBac transposase along with our transposon vector encoding the HA ectodomain and evaluated transposition efficiency by measuring HA surface expression using flow cytometry. The percentage of HA positive cells expressed with the engineered transposase was comparable to that observed with the commercial Super *PiggyBac* transposase, confirming successful integration of the HA construct (Figure 2b). Therefore, the optimized mammalian surface display construct comprised an N-terminal CD5 signal peptide, the HA ectodomain fused to a GCN4 trimerization motif, a C-terminal c-Myc tag, and an IRES–GFP cassette.

### Optimization of transfection and cell surface expression

To determine the optimal conditions for HA surface expression, we tested various amounts of transfected DNA, puromycin for cell selection, and cumate for expression levels (Figure 2e, 2f). The amount of DNA was optimized to balance transfection efficiency with cell viability Transposase and transposon vectors were co-transfected at a 1:2.5 ratio into HEK293T cells seeded in a T25 flask at 80–90% confluency (approximately 1 × 10⁶ cells), with total DNA amounts ranging from 1 to 7.5 µg. At lower DNA concentrations, transfection efficiency and consequently transposition were poor, resulting in minimal HA expression. In contrast, 7.5 µg of total DNA yielded the highest HA binding across the antibody panel, indicating this concentration was optimal for surface display (Figure 2e). DNA concentrations exceeding 7.5 µg were avoided because of reduced cell viability associated with cytotoxic effects from excessive transfection complexes.

Next, we optimized puromycin concentration for the selection of cells with successful HA integration. Concentrations from 1.5 to 6µg/mL were tested. Lower concentrations failed to effectively select for transfected cells only. At 6 µg/mL, the highest percentage of HA-expressing cells was observed, suggesting efficient selection without excessive cytotoxicity (Figure 2f). Puromycin concentrations above 6 µg/mL were avoided because higher antibiotic levels reduced cell viability due to increased cytotoxic stress. Following selection, cumate was added to induce HA expression. Two concentrations, 30 μg/mL and 300 µg/mL were tested. While 30 µg/mL successfully induced HA expression, 300 µg/mL was toxic, leading to complete cell death (Supplementary Figure S9). These results highlight the importance of optimizing the experimental conditions to achieve maximum viral protein surface expression while maintaining cell viability.

To validate the stable integration of HA, HEK293T cells were transfected with either a mixture of transposon and transposase vectors, or the transposon vector alone. A substantial proportion of cells expressed the HA trimer even in the absence of transposase (Figure 2h, Sort 1). To determine whether these HA positive cells had undergone actual transposition, we isolated and expanded them for a second round of sorting. Cells initially transfected with the transposase vector retained HA expression, whereas those transfected without transposase lost positive signal in the second sort, indicating that the initial HA expression was due to transient transfection rather than stable integration (Figure 2h, Sort 2). To further confirm the critical role of *PiggyBac* transposition on HA expression, the HA construct was cloned into a non-*PiggyBac* vector and transiently transfected into HEK293T cells. Comparison of HA positive populations showed that the *PiggyBac* system yielded a higher proportion of HA binders indicating that genomic integration with *PiggyBac* increases HA surface expression (Figure 2g). Based on these optimization studies, all subsequent experiments were performed using 7.5 µg of total DNA for transfection into approximately 1 × 10⁶ HEK293T cells, followed by selection with 6 µg/mL puromycin and induction with 30 µg/mL cumate to achieve robust and stable HA surface expression for downstream analyses.

### Antigenic validation of surface expressed viral proteins

To characterize the folding of surface displayed HA trimers in more detail, we analyzed their antigenic profiles using a panel of monoclonal antibodies known to target diverse HA epitopes by FACS. The panel included antibodies binding to the HA head interface (FluA-20 and S5V2-29), HA stem (429B01), and HA head domain (F045-092). ^35, 36, 38, 39^ Antibody binding was studied across a concentration range from 0.1 nM to 1000 nM. The HA head specific antibody F045-092 showed strong binding across all tested concentrations indicating robust surface exposure of the HA head domain. In contrast, the stem-specific antibody 429B01 showed high binding only at higher concentrations, with minimal binding at lower concentrations, consistent with the more limited accessibility of this epitope on the HA trimer. Similarly, head interface antibodies (FluA-20 and S5V2-29) exhibited robust binding at higher concentrations but reduced binding at lower concentrations, as expected given that their epitope is partially occluded on the pre-fusion HA trimer (Figure 3b).^35^ To validate the antigenicity of surface expressed HA trimers measured by FACS, we performed Biolayer Interferometry (BLI) using recombinantly expressed and purified A/H3N2/Hong Kong/1968 HA trimer against the same antibody panel. The magnitude of the binding responses measured by BLI closely aligned with the FACS binding data, confirming the surface expression of antigenically accurate HA trimers on our platform (Figure 2b).

**Figure 3.**
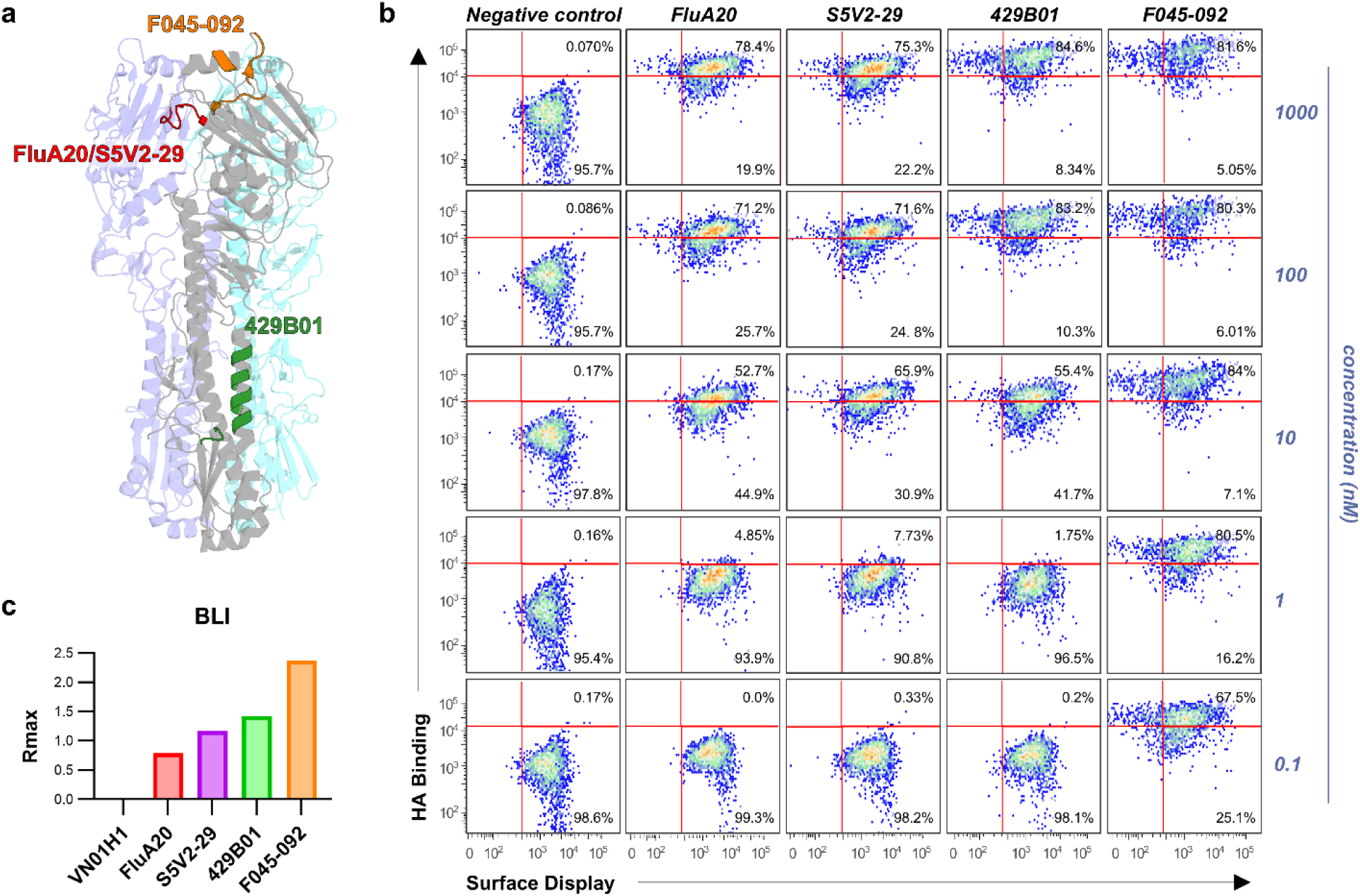
Antigenic profiling of surface displayed A/H3N2/Hong Kong/1968 HA trimers. (a) Structure of the HA trimer highlighting the epitopes recognized by the head-interface antibodies FluA-20 and S5V2-29 (red), the stem-specific antibody 429B01 (green), and the head-specific antibody F045-092 (orange). (b) Flow cytometry analysis of FluA-20, S5V2-29, 429B01 and F045-092 antibody binding to surface-displayed HA trimers. (c) Biolayer interferometry (BLI) analysis of in vitro expressed HA trimer probed with the same antibody panel as in (b), with binding quantified by Rmax values.

To demonstrate that the mammalian display platform could be used for diverse viral proteins, we also characterized the SARS-CoV-2 spike protein and the HIV-1 Env in this system. For SARS-CoV-2, we utilized the 6P-D614G spike derived from the WA-1 strain, which contains six proline substitutions to stabilize the pre-fusion conformation.^42^ We confirmed surface expression of the spike on mammalian cells and studied its antigenic profile using flow cytometry across a panel of monoclonal antibodies. Receptor Binding Domain (RBD) targeting antibodies, B38,^43^ DH1047,^44^ and S309,^45^ exhibited strong binding, indicating robust accessibility of the RBD. The stem helix antibody S2P6^46^showed moderate binding, while the N-terminal domain (NTD) antibody DH1052^44^ demonstrated limited binding, which correlated with their epitope exposure (Figure 4a). These FACS binding data were cross-validated with Enzyme Linked Immunosorbent Assay (ELISA) binding using the recombinant 2P-D614G spike variant.^47^ RBD and NTD antibody binding profiles were consistent across both platforms. However, S2 antibodies showed enhanced binding in ELISA, suggesting that these epitopes are more accessible in the 2P spike variant compared to the more rigid 6P version as previously reported (Figure 4b).^48^

**Figure 4.**
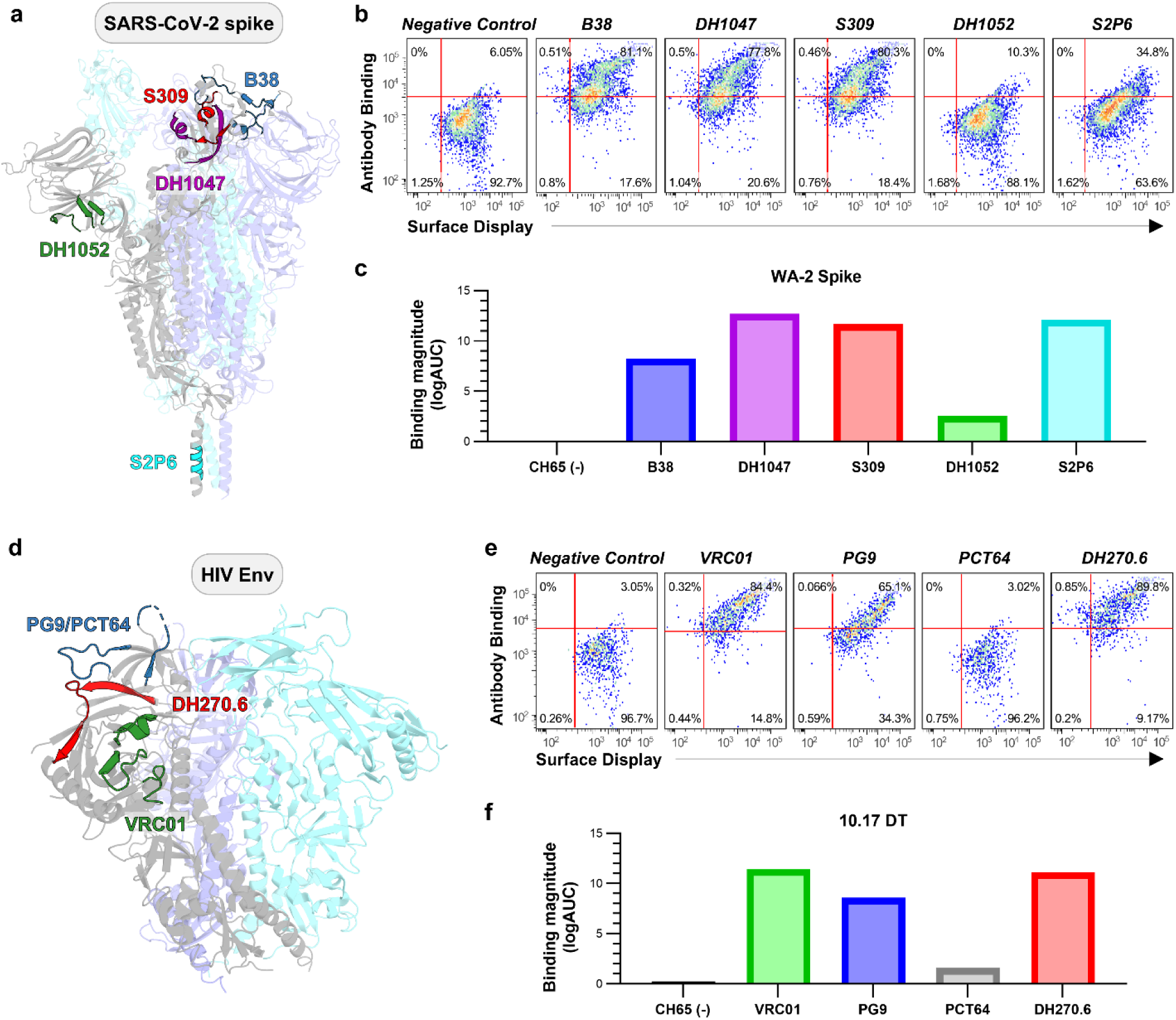
Antigenic profiling of surface-displayed SARS-CoV-2 spike and HIV Env trimers. (a) Structure of the SARS-CoV2 D614G HexaPro trimer highlighting the epitopes recognized by the RBD targeting antibodies B38 (blue), DH1047 (purple), and S309 (red), the N-terminal domain (NTD) antibody DH1052 (green), and the stem helix antibody S2P6 (cyan). (b) Flow cytometry analysis of antibody binding to surface displayed SARS-CoV-2 spikes using antibodies B38, DH1047, S309, DH1052, and the stem helix antibody S2P6. (c) ELISA of recombinant SARS-CoV-2 spike trimers probed with the same antibody panel as in (b). (d) Structural model of the HIV Env 10.17 DT highlighting the epitopes recognized by the CD4 binding site antibody VRC01 (green), the V2 apex antibodies PG9 and PCT64 (blue), and the V3 glycan antibody DH270.6 (red). (e) Flow cytometry analysis of antibody binding to surface displayed HIV Env 10.17 DT trimers using the CD4 binding site antibody VRC01, the V2 apex antibodies PG9 and PCT64, and the V3 glycan antibody DH270.6. (f) ELISA of recombinant 10.17 DT SOSIP trimers probed with the same antibody panel as in (e).

Experimental analysis was also performed on an SOSIP HIV Env trimer, called 10.17DT^9^, to evaluate its surface expression and antigenicity. Flow cytometry was conducted with a panel of monoclonal antibodies targeting the CD4 binding site (VRC01),^49^ the V2-apex (PG9 and PCT64),^50, 51^ and the V3-glycan (DH270.6).^9^ The Env showed high affinity binding to VRC01, PG9, and DH270.6, indicating surface expression and preservation of conformational epitopes during expression. Although PCT64 also targets the V2 apex, its binding was weaker compared to PG9, likely due to differences in epitope recognition mechanisms (Figure 4c). FACS binding results agreed with ELISA binding profiles, confirming the correct antigenic presentation of envelope trimers (Figure 4d). These results show that our mammalian display system can be used to expressing conformationally intact antigens from diverse viruses.

### HA library design and analysis

Next, we developed and screened an HA library to identify mutations that increased binding to HA head trimer antibodies FluA-20 and S5V2-29. This served a dual purpose: 1) to expand the experimental protocol for screening viral protein libraries for desired antigenic properties, and 2) to identify HA mutations with increased affinity to HA head interface antibodies, which are of interest for elicitation as part of a universal influenza vaccine given their broad reactivity across all influenza A strains.

To maintain a tight genotype-to-phenotype linkage in the library, it is critical that the majority of HEK293T transfected cells harbor a single HA library variant. To assess this, we developed an HA displaying plasmids that co-expressed mCherry. Co-transfection of the two HA plasmids containing GFP or mCherry in equal amounts (7.5 µg total DNA) revealed that >30% of the cells expressed both fluorescent reporters, indicating the presence of different plasmid variants in the same cell. To minimize this effect, a carrier transposon plasmid lacking the HA insert and without resistance to puromycin was added during transfection to dilute the amount of target DNA.^19^ The empty transposon vector contains the same ITR sequences as the HA transposon vectors but lacks an HA insert, allowing it to integrate into the genome as a carrier. Co-transfection with this carrier vector reduces the likelihood that individual cells acquire multiple copies of the GFP- and mCherry-tagged HA constructs during transposition. GFP and mCherry HA vectors were diluted 10-fold with the carrier vector, keeping the total DNA amount constant (7.5µg). This adjustment led to a modest decrease in HA expression but led to an approximately 75% decrease in the proportion of GFP–mCherry double-positive cells. (Figure 2j). Following transposition, puromycin selection removes cells that carry only the carrier vector, thereby enriching for puromycin resistant cells that maintain stable HA expression with predominantly single-copy integration.

Once optimal transformation conditions have been established, we designed a single-site saturation library at HA head sites located within 10Å of the 220-loop, the epitope of HA head-trimer antibodies such as FluA-20 and S5V2-29 (Figure 5a). The 220-loop is completely occluded in the pre-fusion HA structure, although it must become exposed through molecular “breathing”, since antibodies against this site bind the recombinant protein as shown above (Figure 3a) and protect against live virus challenge.^35, 36^ The HA library consisted of all possible amino acid variations at 110 sites that may affect epitope exposure and improve antibody binding, either directly or allosterically, yielding a total of 2090 HA variants.

**Figure 5.**
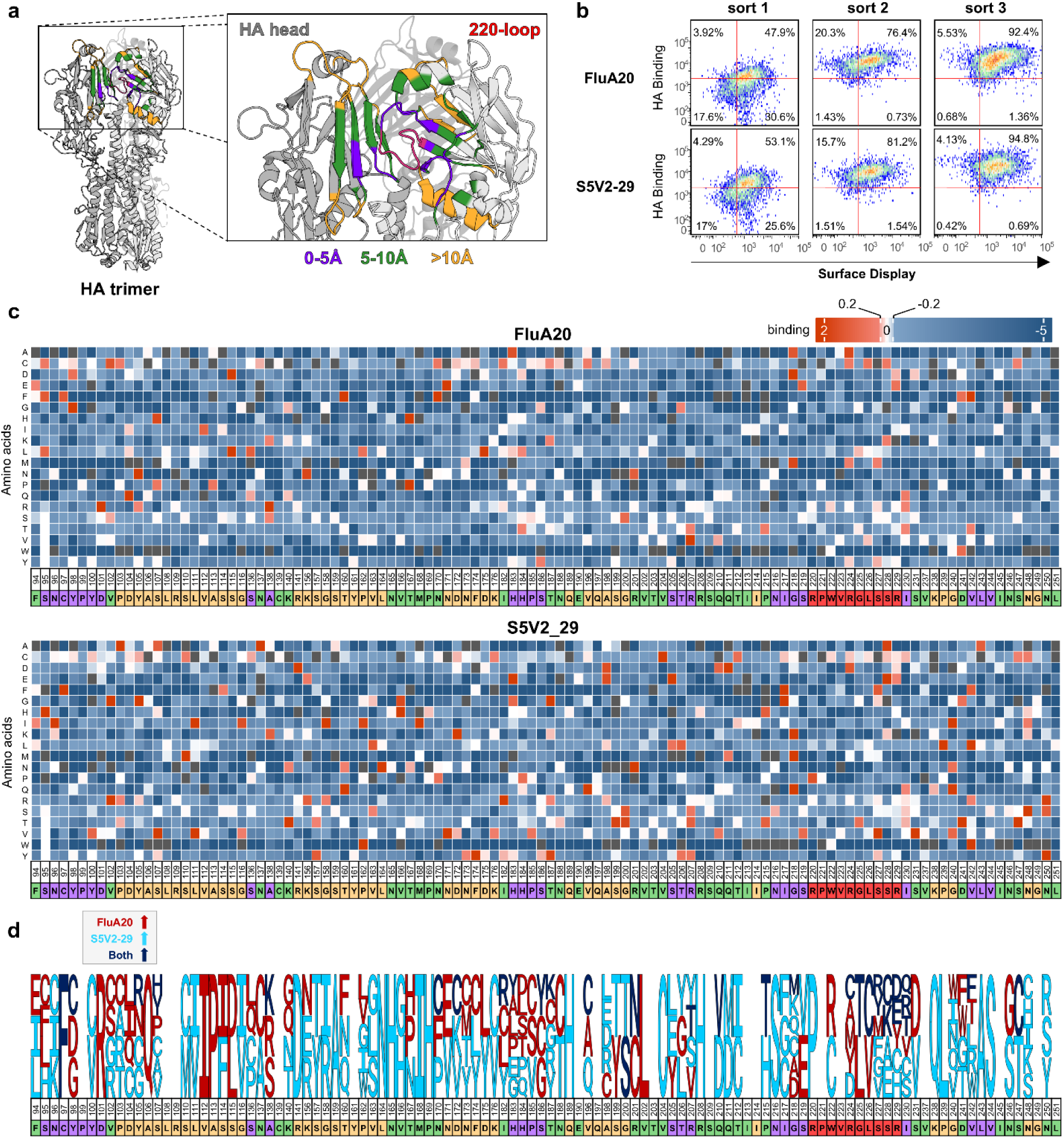
Analysis of HA head mutant libraries using the mammalian surface display platform. (a) Structural model of the HA trimer with a detailed view of the regions sampled in the single site saturation library. The 220-loop is highlighted in red, residues within 5 Å of the 220-loop are shown in purple, residues between 5-10 Å are shown in green, and residues located beyond 10 Å are shown in orange (b) Flow cytometry analysis showing enrichment of HA variants binding the head interface antibodies FluA20 and S5V2-29 over three iterative rounds of antibody-based sorting. (c) Heatmaps showing the frequency of amino acid substitutions across the HA head region following antibody-based selection with FluA-20 and S5V2-29 after three rounds of library selection. Effect of every mutation on binding to each antibody calculated as enrichment levels (blue: unfavorable; red: favorable; white: neutral) excluding the wild type for each position and the mutations with frequencies less than 0.01% in the naive library (dark grey). Residues are colored according to their spatial grouping relative to the 220-loop, as defined in panel (a). (d) Enriched amino acid substitutions across the HA head region following antibody-based selection. Mutations enriched in FluA-20 selected populations are shown in red, mutations enriched in S5V2-29 selected populations are shown in cyan, and mutations enriched under both selection conditions are shown in blue

To identify HA mutations that improve recognition by head interface antibodies, the HA mutant libraries were subjected to antibody selection by FACS using FluA-20 or S5V2-29 over three iterative rounds of sorting. At each round, cells binding to the target antibodies were collected and expanded. The second round showed the highest increase in the proportion of antibody binders, with the third round further enriching for cells that bound FluA20 or S5V2-29 (Figure 5b).

Next Generation Sequencing (NGS) was performed on the initial, unsorted HA library and on cells collected following the second and third selection rounds. For a given mutation, its frequency was compared in the unsorted library and at each selection round; mutations that increased in frequency upon antibody selection were considered favorable for FluA-20 or S5V2-29 binding, while those that were depleted upon sorting were considered unfavorable. As expected, most of the mutations in the 220-loop were detrimental to antibody binding. This observation is consistent with prior mutational analyses showing that substitutions at 220-loop residues such as Val223 and Arg229 substantially reduce FluA-20 binding,^36^ supporting the validity of our NGS-based selection results (Figure 5c). Selection with FluA-20 yielded a more limited set of enriched mutations compared to S5V2-29, which may reflect more stringent structural or conformational requirements for FluA-20 recognition, whereas S5V2-29 appears to accommodate a broader range of sequence changes that enhance epitope exposure. (Figure 5c). Of the 60 sites where mutations favorable for FluA-20 binding were identified, 16 were located within 5Å of the epitope, 24 between 5Å and 10Å and 22 further away than 10Å. Similarly, for S5V2-29, out of 87 sites where mutations favorable for S5V2-29 binding were identified, 23 were located within 5Å of the epitope, 23 between 5Å and 10Å and 33 further away than 10Å. Comparison of clones selected with FluA-20 and S5V2-29 identified only a limited set of mutations enriched under both selection conditions. Among the mutations, 9 were located within 5Å of the epitope, 6 between 5–10Å, and 5 at distances greater than 10Å, indicating that residues outside the core epitope can contribute to antibody recognition through indirect or allosteric effects (Figure 5d).

## Discussion

The development of a mammalian surface display platform using the *PiggyBac* transposon system provides a robust and versatile framework for engineering complex viral glycoproteins. By enabling stable genomic integration and inducible expression of viral proteins, our display platform addresses key limitations in existing transient and lentiviral mammalian display systems. Transient transfection approaches often suffer from heterogenous expression, batch to batch variability and loss of expression after each cycle of transfections.^52, 53^ The lentiviral systems require time intensive virus packaging and pose biosafety concerns.^54-56^ The *PiggyBac* transposon system provides an efficient and scalable alternative for integrating large transgenes^34, 57-59^ in mammalian cells, making it suitable for high throughput library expression of viral antigens.

Our platform combined the *PiggyBac* integration machinery with a PGDFR transmembrane anchoring protein and a cumate inducible switch that enables controlled surface expression of viral proteins such as influenza HA, HIV Env, and SARS-CoV-2 spike. Stable expression of correctly folded trimeric proteins on the mammalian cell surface was validated by antibody binding, and these findings were consistent with *in vitro* biophysical characterization using conformation-sensitive antibodies (Figures 3 and 4). Incorporation of trimerization and reporter elements, such as the GCN4 motif and an IRES-linked GFP, further improved stability and surface expression (Figure 2c and 2d). Importantly, these optimizations significantly improved expression without compromising antigenicity as validated across diverse viral proteins.

From a methodological standpoint, the cost-effective use of engineered hyperactive *PiggyBac* transposase represents a practical advantage over commercial transposase systems which are only single use. We demonstrated that the non-commercial transposase performs comparably to commercial plasmid supporting its use in high throughput settings (Figure 2b). Moreover, the inclusion of the cumate-inducible promoter enables precise control over protein expression levels, preventing cellular stress during stable cell line generation and facilitating the expression of complex viral proteins on the cell surface. One of the major technical challenges in mammalian display systems is achieving stable, single copy integration to maintain a clear genotype-phenotype linkage for library screening. Our results demonstrate that *PiggyBac* transposition enables single copy integration through DNA co-transfection strategies (Figure 2i). This feature is critical for downstream analyses, ensuring that each cell displays a unique library variant for accurate mapping of antigenic landscapes.

Using this system, we successfully expressed and characterized a library of HA head variants designed to enhance the accessibility of the conserved HA head interface epitope, which is targeted by broadly neutralizing antibodies such as FluA-20 and S5V2-29. While early selection rounds yielded variants with modest improvements in binding, further rounds under more stringent selection conditions may enrich for higher affinity mutants. These results validate the feasibility of our approach for iterative oligomeric antigen engineering in mammalian cells. Based on NGS analysis, we identified mutations that enhance binding to both FluA-20 and S5V2-29, with these mutations distributed across the HA head region (Figure 5d). Future studies will focus on expressing individual HA mutants to enable detailed biophysical characterization and evaluation in immunization studies.

Beyond influenza HA our findings with SARS-CoV-2 spike and HIV Env indicate that this platform can accommodate diverse viral glycoproteins requiring proper folding and mammalian glycosylation for native antigenicity. The spike and Env proteins expressed using our surface display exhibited expected binding profiles to epitope-specific monoclonal antibodies targeting RBD, stem, and glycan epitopes (Figure 4a, 4c). These findings highlight the that the display platform maintains the conformational integrity of structurally complex viral proteins.

The mammalian display platform described here offers new opportunities for immunogen design. It combines the high throughput capability of microbial display systems with the physiological relevance of mammalian expression. Unlike yeast or phage display, this system allows direct screening of proteins with native glycosylation and trimeric structures enabling accurate evaluation of conformational epitopes recognized by neutralizing antibodies. The combination of stability, scalability, fidelity, and safety makes this approach well suited for developing optimized immunogens for universal vaccines. This platform provides a practical framework for advancing engineered viral antigens toward optimized vaccine candidates through iterative and scalable antigen design.

## Supplementary Figure Legends

**Supplementary Figure 1.**
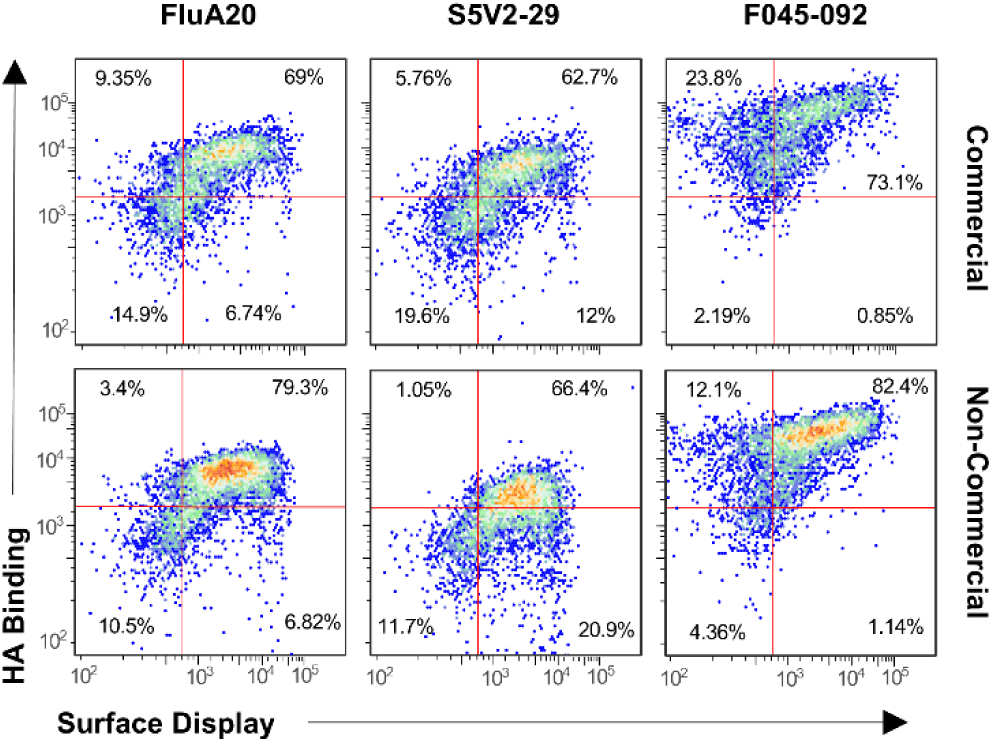
Flow cytometry analysis comparing HA surface expression using a commercial PiggyBac transposase and a non-commercial engineered PiggyBac transposase.

**Supplementary Figure 2.**
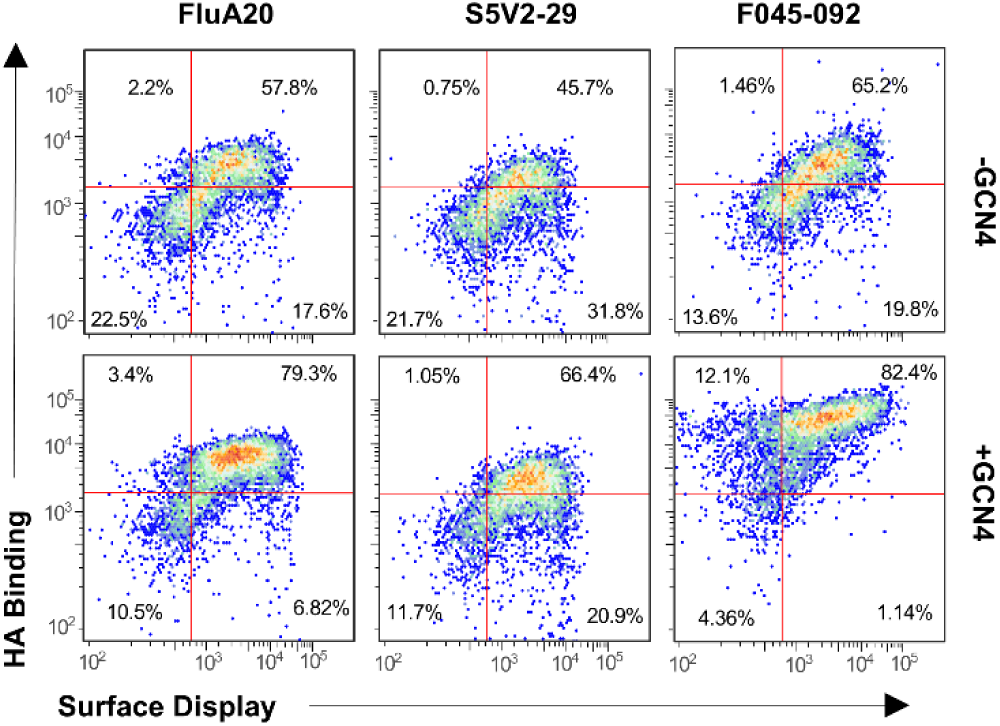
Flow cytometry analysis comparing HA surface expression in constructs expressed with or without the GCN4 trimerization motif.

**Supplementary Figure 3.**
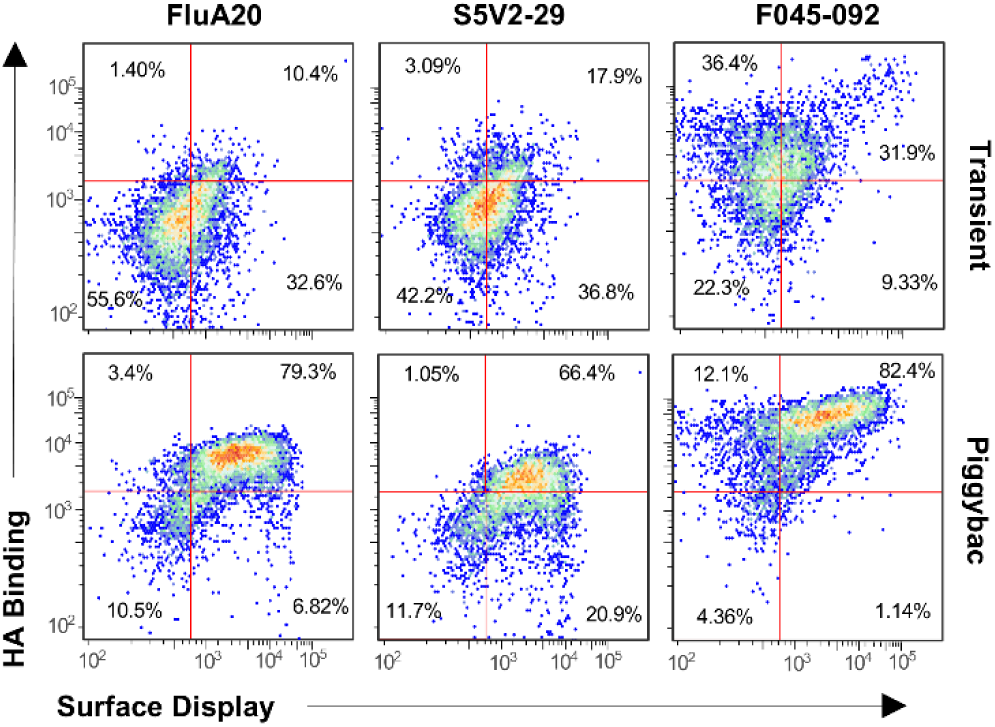
HA surface expression following PiggyBac mediated genomic integration versus transient transfection.

**Supplementary Figure 4.**
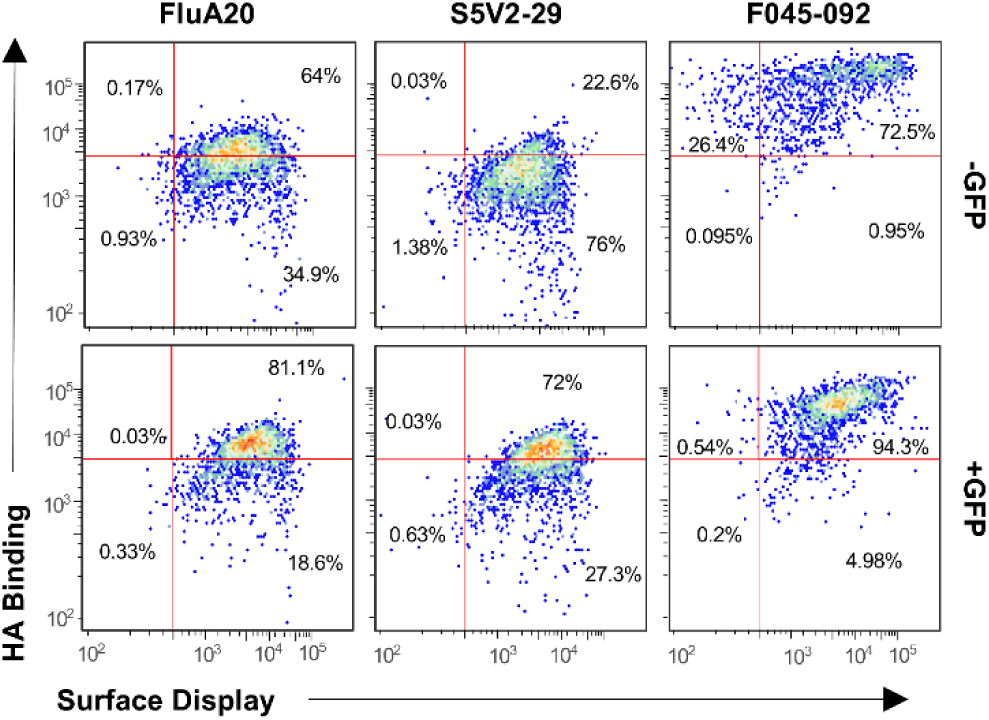
Flow cytometry analysis comparing HA surface expression in constructs lacking or containing an IRES–GFP expression cassette.

**Supplementary Figure 5.**
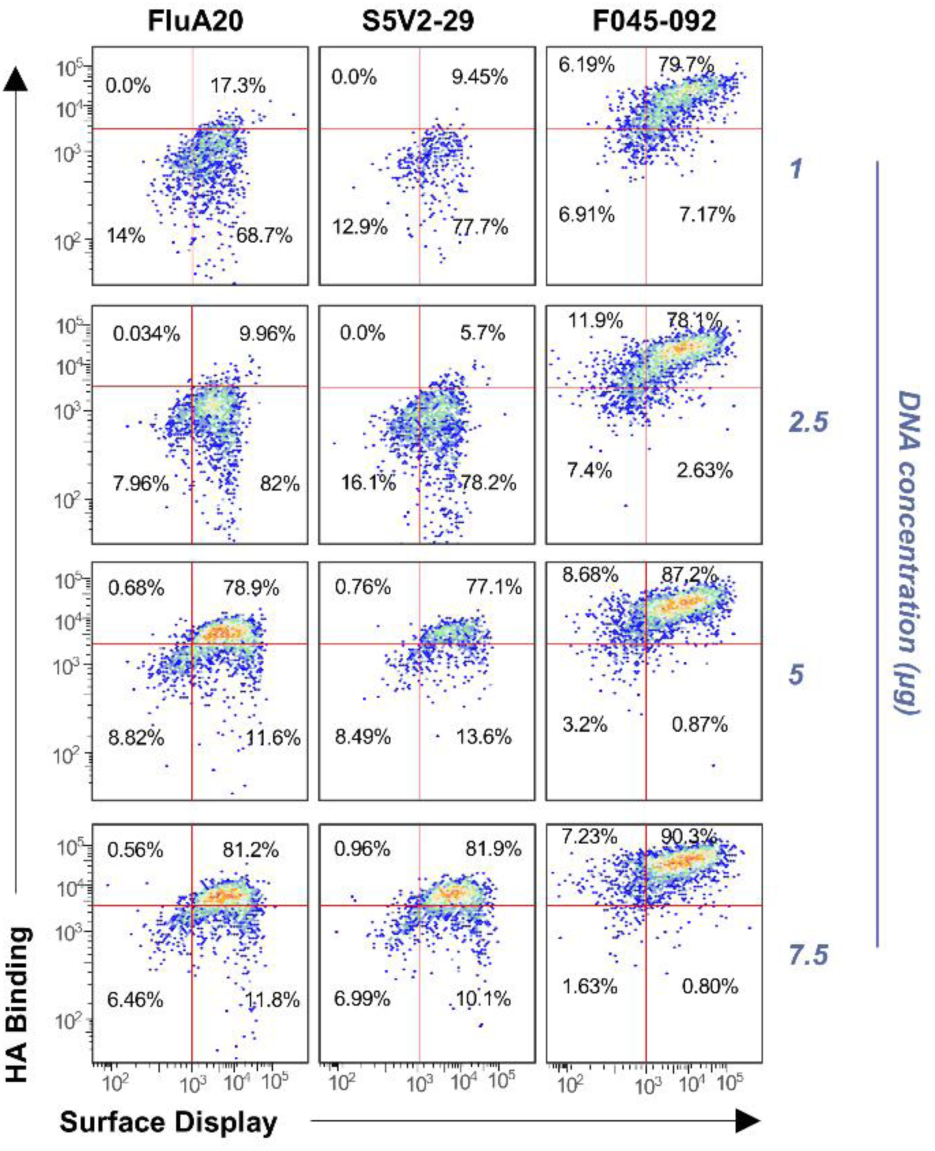
Flow cytometry analysis showing the effect of varying total DNA amounts on HA surface expression.

**Supplementary Figure 6.**
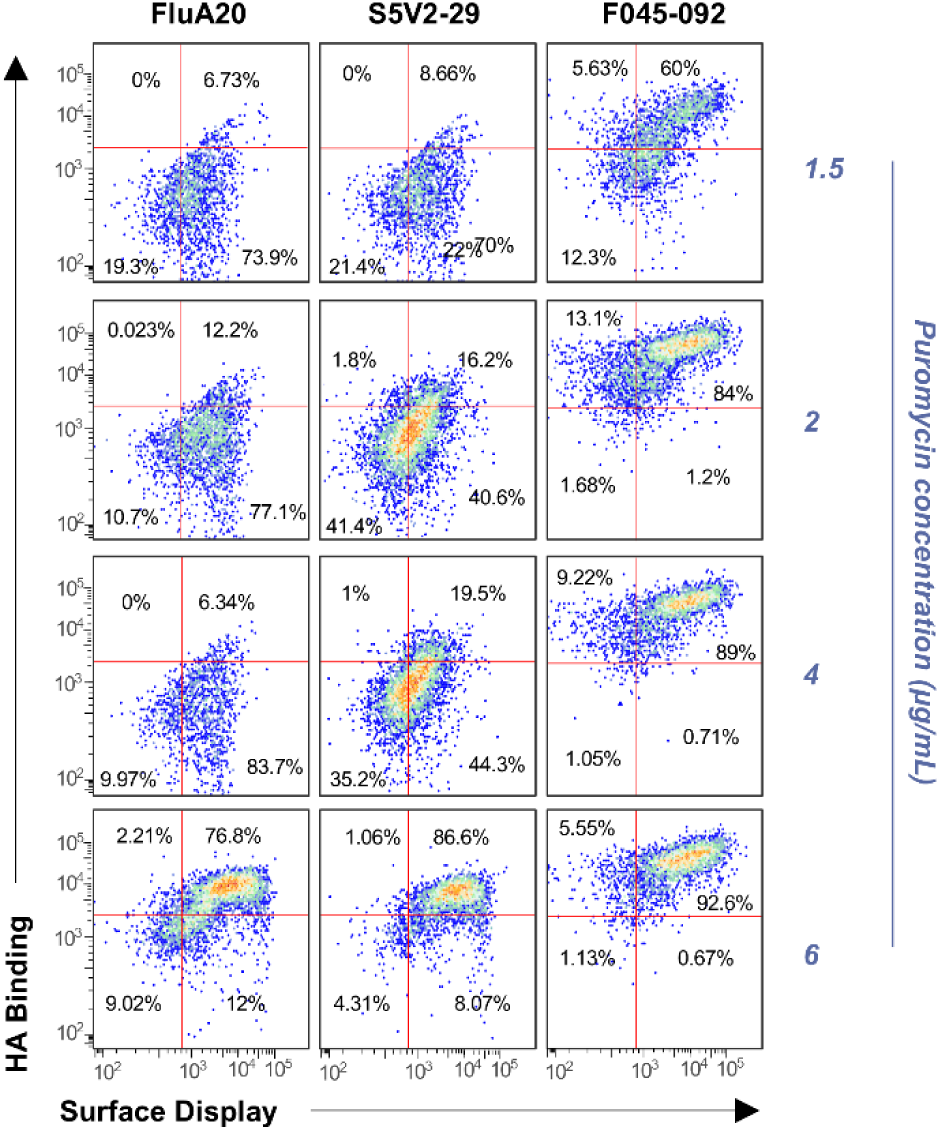
Flow cytometry analysis showing the effect of varying puromycin concentrations on selecting for HA expressing cells.

**Supplementary Figure 7.**
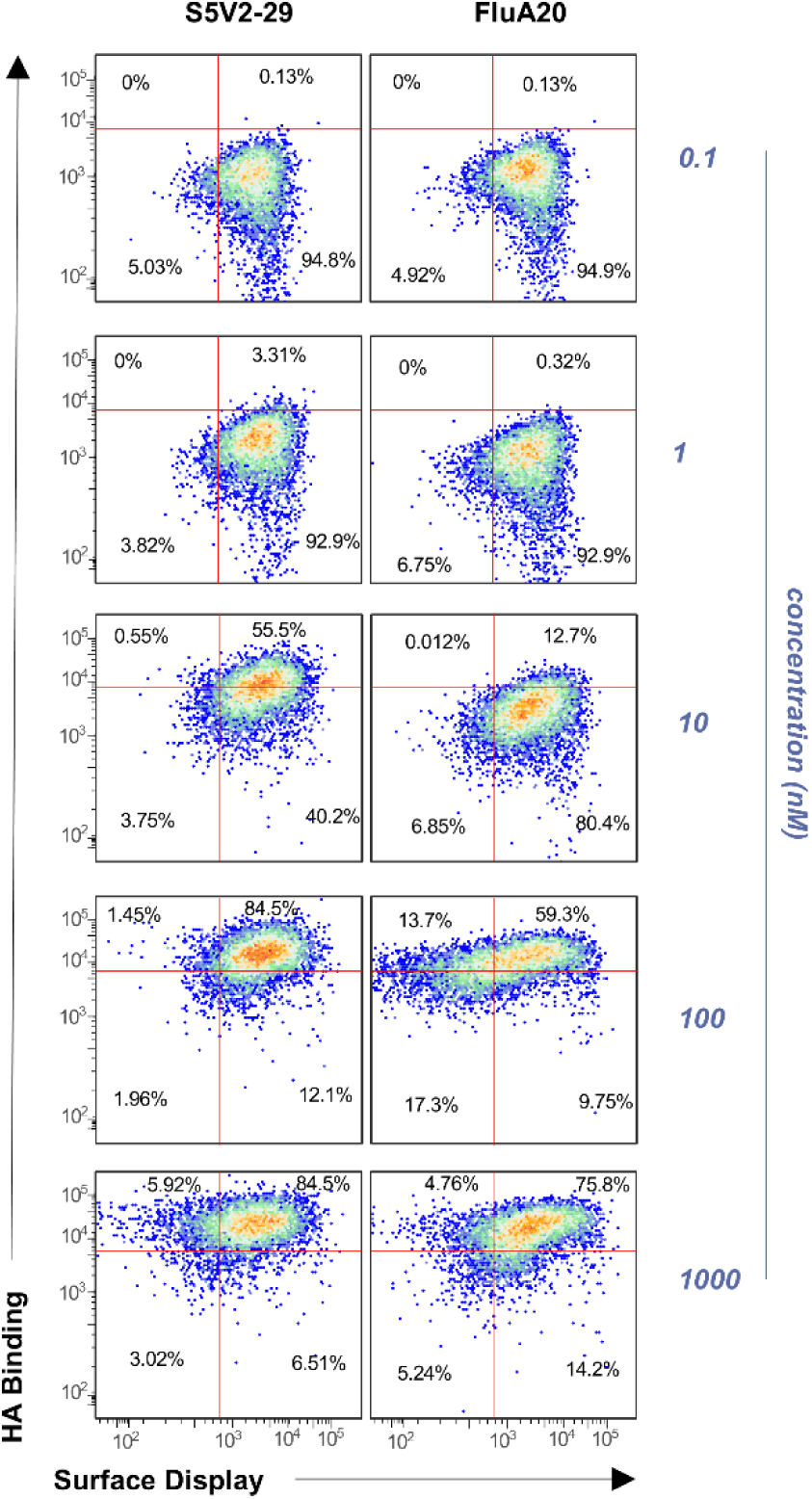
Antibody titration analysis of the enriched HA libraries following the second round of sorting, showing concentration dependent binding to the head-interface antibodies FluA20 and S5V2-29.

**Supplementary Figure 8.**
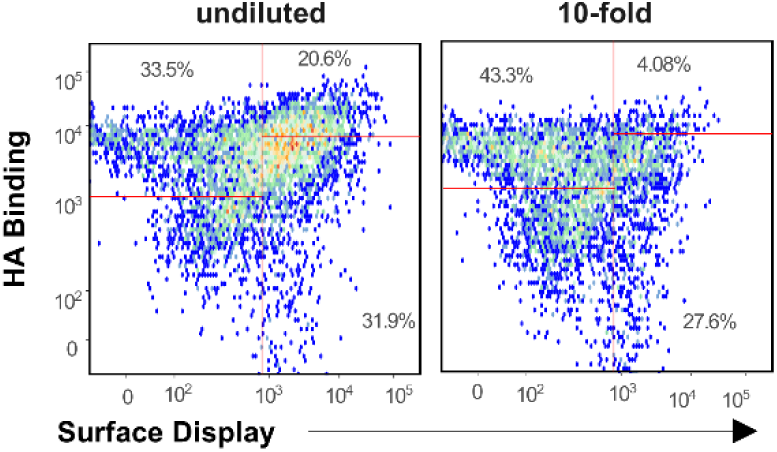
Flow cytometry analysis of GFP and mCherry expression showing reduced GFP+ mCherry double positive populations following dilution with the carrier transposon.

**Supplementary Figure 9.**
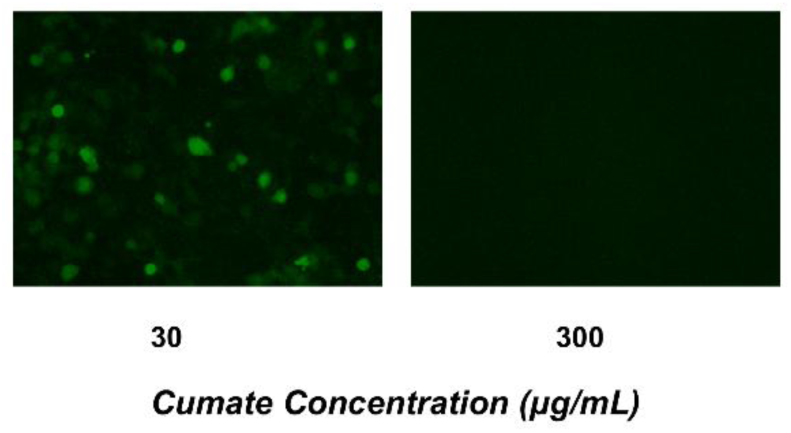
Fluorescence microscopy images showing HA and -GFP co0-expression following induction with 30 µg/mL cumate and complete cell death following induction with 300 µg/mL cumate.

## Materials and Methods

### Plasmids, Strains, Media and Chemicals

Inducible PiggyBac transposon cloning vectors used were PB-CuO-CMV-MCS-EF1a-CymR-T2A-Puro (PBQM800A-1, System Bioscience), PB-Cuo-shMCS-IRES-GFP-EF1α-CymR-Puro (PBQMSH812A-1, System Bioscience) and PB-EF1α-MCS-IRES-Neo (PB533A-2, System Biosciences). The transposase vectors used were Super piggyBac Transposase expression vector (PB200A-1, System Bioscience) and hyperactive piggyBac transposase (pBase) described in Yusa et.al ^41^. All the cloning was done in NEB 5-alpha competent *E. coli* cells (New England Biolabs). Transformed *E. coli* cells were grown in Luria–Bertani broth (LB, Apex Bioresearch Products) supplemented with 100 μg/mL ampicillin. Hek293T cells (ThermoFisher Scientific) were used for expression of mammalian display constructs in Dulbecco’s Modified Eagle’s Medium (DMEM, ATCC). Transfection Reagent VirusGen (MirusBio) was used for all the transfections, 10 mg/mL Puromycin (ThermoFisher Scientific) was used for selection. Protein expression on the surface was induced by Cumate solution 10,000 X (QM100A-1). For fluorescent labeling FITC conjugated chicken anti-cmyc antibody (ICL Labs) and R-Phycoerythrin AffiniPure^®^ Goat Anti-Human IgG, Fcγ fragment specific (Jackson ImmunoResearch) were used. Unless mentioned otherwise, mammalian surface display plasmids were constructed by Genscript. The HA head libraries were synthesized by Twist Bioscience and all oligo DNA were synthesized by Integrated DNA Technologies.

### Plasmid Construction

The mammalian display DNA constructs were synthesized by Genscript, and included H3/Hong Kong 1968 ectodomain, cmyc tag, GCN4 trimerization motif, and PGDFR membrane domain. The PiggyBac transposon vectors PB-CuO-CMV-MCS-EF1a-CymR-T2A-Puro and PB-Cuo-shMCS-IRES-GFP-EF1α-CymR-Puro were cut with Not I and Nhe I (New England Biolabs) at the multiple cloning sites. The HA ectodomain insert from Genscript was amplified with PCR using Q5 High Fidelity 2X master mix (New England Biolabs) using the manufacturer’s protocol. The inserts were gel purified and cloned into the cut vectors of PB-CuO-CMV-MCS-EF1a-CymR-T2A-Puro and PB-Cuo-shMCS-IRES-GFP-EF1α-CymR-Puro with T4 DNA Ligase using the manufacture’s protocol. The cloned plasmids were transformed in NEB 5-alpha *E. coli* cells and the cloned plasmids were isolated with Zymopure Plasmid Miniprep kit (Zymo Research) and verified by Sanger Sequencing (Genwiz).

### PiggyBac Transfection, Selection, and Induction

Transposase and transposon plasmids (7.5 μg total DNA in ∼2 x 10^6^) were transfected on 80% confluent HEK293T cells using Transfection Reagent VirusGen (MirusBio) at a ratio of 1:2.5 μg transposase: transposon. The DNA-transfection reagent mix was buffered for 20 minutes in Gibco OptiMem (ThermoFisher Scientific). The cell culture media was replaced with new media before adding the transfection mixture. Transfected cells were incubated at 37°C. Three days post transfection, cell culture media was replaced with DMEM supplemented with 6 μg/mL Puromycin and incubated at 37°C. After 3 days of puromycin treatment, the media was changed to DMEM supplemented with 1X cumate (30 μg/mL). Cells were subjected to induction for three days before being analyzed by flow cytometry. Using the same protocol, total plasmid DNA concentrations of 1, 2.5, 5, puromycin concentrations of 1.5, 2, 4 and cumate concentration of 300 μg/mL were tested for the optimization of protocol.

### Antibody Production

Antibodies were produced in Expi293F cells (ThermoFisher Scientific). The cells were transiently transfected at a density of 2.5 × 10^6^ cells/mL with an equimolar plasmid mixture of heavy and light chains using Expifectamine (Invitrogen). Five days post-transfection, the cells were separated from culture media by centrifugation, and the supernatant was filtered with a 0.8 µm filter. The supernatant was incubated with equilibrated Protein A beads (ThermoFisher) for 1 hour at 4°C. Beads were washed with 20 mM Tris, 350 mM NaCl at pH 7, and antibodies were eluted with a 2.5% Glacial Acetic Acid elution buffer, and buffer was exchanged into 25 mM Citric Acid, 125 mM NaCl buffer at pH 6. Antibody expression and purity were confirmed by SDS-PAGE analysis and quantified by measuring absorbance at 280 nm (Nanodrop 2000).

### Testing antigenicity with flow cytometry

The induced cells were collected and harvested at 125 *g* for 5 minutes at 4°C. The harvested cells were washed with 1X Phosphate Buffered Saline (PBS, Thermofisher Scientific) supplemented with Bovine Serum Albumin (BSA, Sigma Aldrich). The cells were resuspended in primary labeling reagent (1:10 Flu antibody: buffer) and incubated on ice for 1 hour. Cells were then resuspended in secondary labeling reagent (1:20 anti cmyc FITC: buffer and 1:20 anti human IgG: buffer) and incubated on ice for 30 minutes. The cells were washed, resuspended in the buffer, and analyzed for double positives binding to FITC and PE using the SONY SH800 sorter.

### Imaging with fluorescence microscope

The transposon and transposase plasmids, PB-Cuo-shMCS-IRES-GFP-EF1α-CymR-Puro-HA, PB-Cuo-shMCS-IRES-mCherry-EF1α-CymR-Puro-HA (Genscript), PB-EF1α-MCS-IRES-Neo and pBase were transfected into Hek293T cells as we described previously. (The ratio of transposons GFP: mCherry: Neo= 0.5: 0.5:9). The cells with positive Piggybac integration were selected with Puromycin as mentioned in the previous methods. Three days post puromycin selection cells were transferred to a Cell Culture Microplate, 96 well clear bottom black plate (Griener) coated with Poly-L-Lysine. The cells were treated with 1X Cumate and incubated at 37 °C for three days. Before imaging the cells, the media was replaced with 1% Gibco HEPES buffer solution (ThermoFisher), 1% FBS and Hanks’ Salt Solution (HBBS 1X, ThermoFisher). Cells were imaged in 96 well plates with imaging clear plastic bottoms (Greiner Bio) using a Nikon 20X Lambda D, 0.8 NA air objective on Nikon Eclipse Ti2 inverted microscope and using Elements acquisition software and LED illumination (Lumencor). Excitation filters: dichroic mirrors: emission filters used were - ET555/20x: T585lxpr: ET595/33m for mCherry; ET480/30x: 89402bs: ET519/26m for GFP-all filters and dichroics Chroma technology unless noted. Images were obtained using an Orca Fusion sCMOS camera (Hamamatsu).

### Binding studies using biolayer interferometry (BLI)

A Sartorius Octet R4 instrument was used to measure binding of H3/Hong Kong 1968 to a panel of antibodies (VNO1H1, FluA-20, S5V2-29, 429B01 and F045-092). All the assays were performed in 1X HBS-EP Buffer (Cytiva) supplemented with 0.1% BSA. The final volume for all the solutions was 200 μl per well. Assays were performed at 25°C in solid black tilted-bottom 96-well plates (Griener). Human antibodies (2.5 μg/ ml) in 1X HBS-EP buffer was used to load Octet ProA Biosensors (Sartorius) for 90 seconds. Biosensor tips were then equilibrated for 60 seconds in 1X HBS-EP,0. 1% BSA buffer before binding assessment. Binding of HA trimer was allowed for 210 seconds, and dissociation was performed for 420 seconds. Data analyses were carried out using Octet Analysis Studio software, version 13.0.

### Antibody Binding by Enzyme-linked immunosorbent assays (ELISA)

Antigens were coated overnight on 384-well plates (Corning, 3700) at 4°C. Purified antibodies were used at 100 µg/mL starting concentrations and were then serially diluted 3-fold and incubated at room temperature for 1 hour. Goat anti-human IgG-HRP secondary antibody (Jackson ImmunoResearch Laboratories, 109-035-098) was diluted 1:15,000 and added to plates, incubated for 1 h, and developed using TMB substrate (SureBlue Reserve, KPL, 5120-0083). Plates were read at 450 nm on a Cytation 1 plate reader (BioTek). Data were analyzed and plotted using GraphPad Prism version 10.2.0 for Windows (GraphPad Software, Boston, Massachusetts USA, www.graphpad.com). Statistics were performed by the Wilcoxon signed-rank test.

### HA head library design

For the HA head single site variant library, one hundred and ten residues, either within 10A of the 220-loop, or connected to these residues though secondary structures, were singly randomized to all possible combinations of amino acids each site, giving a total library size of 2090 variants.

### HA head library assembly

The HA head libraries were synthesized by Twist Biosciences. Libraries were PCR amplified with Q5 High Fidelity Polymerase and purified. The purified library insert was cloned into PB-Cuo-shMCS-IRES-GFP-EF1α-CymR-Puro cut vector at NheI and SwaI restriction sites using 2X Gibson Assembly Master Mix (New England Biolabs), according to the manufacturer’s instructions. The cloning was confirmed by Sanger Sequencing. The library was then transformed in NEB 5-alpha cells and the DNA was extracted with Pure yield plasmid midiprep system (Promega). The HA head variant library expressed in E. coli yielded >10000 clones. The DNA pool was extracted and transfected into HEK293T cells.

### HA head library sorting

PB-Cuo-shMCS-IRES-GFP-EF1α-CymR-Puro-HA library, PB-EF1α-MCS-IRES-Neo and pBase were transfected into Hek293T (The ratio of transposons HA library: Neo= 1:9), puromycin selection and cumate induction were done as we described before. Three days post induction cells were harvested and labeled according to previously described protocol with FluA-20 and S5V2-29 antibodies. Cells were sorted on a SONY SH800 sorter. Approximately 5 × 10^4^ double positive cells were collected and expanded for approximately one week and induced with 1X Cumate. A second round of sorting was performed three days post induction.

### NGS

Once the desired population had been obtained, chromosomal DNA was extracted from the cell culture with the GenElute Mammalian Genomic DNA Miniprep Kit (Sigma). The HA library gene was PCR amplified from the genomic DNA and inserted back into pCDNA3 vector via Gibson Assembly and transformed into NEB 5-alpha competent cells and colonies were sequenced at Genewiz. DNA samples were prepped and run using the Illumina MiSeq v3 reagent kit following manufacturer’s protocols. Illumina sequencing returned an average of 48.9 million reads per sample, of which an average of 31 million mapped to the HA head amplicon. Sequencing data was processed using Geneious Prime to compute the amino acid frequency and distribution.

## Author Contributions

J.C. and M.L.A. designed the study. Experimental work was done by J.C., C.H., B.R., and D.M. Data analysis was performed by J.C., B.R., and M.F. The study was directed by M.L.A. The manuscript was written by J.C. and M.L.A (first draft) with input from the other authors.

## Acknowledgments

We would like to acknowledge the Duke Human Vaccine Institute Genomics Core for assistance with next-generation sequencing. This work was supported by the National Institutes of Health (NIH) through grants R01AI168337 (M.L.A).

## Notes

### Competing Interest Statement

The authors have declared no competing interest.

### Summary of Updates

The revision was done to change the authors' positions.

